# The Structure-Selective Endonucleases GEN1 and MUS81 are Functionally Complementary in Safeguarding the Genome of Proliferating B Lymphocytes

**DOI:** 10.1101/2022.02.09.479711

**Authors:** Keith Conrad Fernandez, Laura Feeney, Ryan M. Smolkin, Wei-Feng Yen, Allysia J. Matthews, John H. J. Petrini, Jayanta Chaudhuri

## Abstract

During the development of humoral immunity, activated B lymphocytes undergo vigorous proliferative, transcriptional, metabolic, and DNA remodeling activities; hence, their genomes are constantly exposed to an onslaught of genotoxic agents and processes. Recombination-dependent DNA transactions that preserve the integrity of the genome and of the DNA replication process generates Holliday junctions that must be eliminated for the accurate segregation of sister chromatids and faithful propagation of genomic material. To investigate the role of two Holliday junction resolvases, GEN1 and MUS81, in B cell biology, we established B-cell conditional knockout mouse models and found that targeted deletion of GEN1 and MUS81 in early B cell precursors halts their development and maturation while selective loss of the resolvases in mature B cells inhibits the generation of robust germinal centers. Upon activation, these double-null mature B lymphocytes fail to proliferate and survive while exhibiting transcriptional signatures of p53 signaling, apoptosis, and type I interferon response. Metaphase spreads of these resolvase-deficient cells showed severe and diverse chromosomal abnormalities, including a preponderance of chromosome breaks, consistent with a defect in resolving DNA recombination intermediates. These observations underscore the essential roles of GEN1 and MUS81 in safeguarding the genome to ensure the proper development and maintenance of B lymphocytes.

## INTRODUCTION

B lymphocytes comprise the humoral arm of the adaptive immune system. They undergo a well-orchestrated series of clonal expansion and differentiation programs in the bone marrow to become mature B cells that reside in secondary lymphoid organs such as the spleen and lymph nodes (LeBien and Tedder, 2008; Pieper et al., 2013). During an adaptive immune response, B cells are recruited into the germinal center (GC) where they adopt one of two cellular fates before exiting the GC: memory B cells that confer immunological memory and plasma cells that produce antibodies of high affinity and specificity (Victora and Nussenzweig, 2012). B lymphocytes are unique among other immune cells in that they initiate programmed DSBs both as developing precursors in the bone marrow and as GC B cells in the secondary lymphoid organs (Alt et al., 2013). V(D)J recombination generates a primary repertoire of B cell receptors that can be further diversified when GC B cells employ the DNA-modifying enzyme activation-induced cytidine deaminase (AID) to instigate the formation of DSBs in the immunoglobulin heavy loci (IgH) for class-switch recombination (CSR) and to somatically hypermutate the variable regions of the immunoglobulin loci (Feng et al., 2020; Schatz and Swanson, 2011; Xu et al., 2012). To outcompete other clonal cells and be selected for survival and differentiation, GC B cells not only are required to express the correct antigen-specific receptor of high affinity and specificity, but they must also satisfy the formidable replicative, transcriptional, and metabolic demands for clonal expansion (Young and Brink, 2021). These cellular activities pose significant collateral genotoxic hazards to GC B cells and thus, safeguarding the cells’ genomic integrity is paramount for the faithful duplication and propagation of genetic information.

Impediments to the progression of the replication fork—termed replication stress— represent a significant endogenous source of DSBs in proliferating cells, producing up to 50 DSBs per cell cycle in a cell (Mehta and Haber, 2014; Zeman and Cimprich, 2014). Factors that impair the functionality of the replication machinery include repetitive sequences, secondary structures (such as R-loops and G-quadruplexes), transcription-replication conflicts, lesions such as thymidine dimers and single-stranded breaks, oxidative stress, and imbalance or depletion of the nucleotide pool (Zeman and Cimprich, 2014). To ensure completion of DNA synthesis before mitosis commences, replication forks that have stalled or collapsed can be restarted via recombination-dependent or - independent pathways, the choice of which is contingent upon the nature of the replication barrier, the duration of stalling, the nature of the intermediates generated after fork stalling, and the type of processing these intermediates undergo (Berti et al., 2020; Petermann and Helleday, 2010; Zeman and Cimprich, 2014). Though primarily studied in the context of DSB repair, proteins involved in homologous recombination (HR) including BRCA2 and RAD51 are also critical for the protection, remodeling, and recombination-dependent restart of replication forks, highlighting the necessity of HR in mitigating replication stress and assisting timely completion of DNA synthesis (Ait Saada et al., 2018; Carr and Lambert, 2013; Scully et al., 2021).

Recombination-dependent repair of DSBs and restart of replication forks entail strand invasion and homology search of an intact duplex DNA, culminating in the formation of Holliday junctions (HJs) that physically link the two sister chromatids (Falquet and Rass, 2019). Several mechanisms have evolved to process these intermediates, as failure to eliminate HJs results in the entanglement of sister chromatids that precludes chromosomal disjunction and propagation of the correct complement of genetic material (West and Chan, 2017). The BLM-TOP3A-RMI1-RMI2 (BTR) complex dissolves double HJs to generate non-crossover products while structure-selective endonucleases (SSEs) such as MUS81 (in complex with EME1, SLX1-SLX4, and XPF-ERCC1) and GEN1 resolve single and double HJs to generate both crossover and non-crossover products, depending on the position of the nicks introduced (Blanco and Matos, 2015). Because these SSEs are active against a broad spectrum of branched DNA structures, they are subjected to multiple cell cycle-dependent regulatory mechanisms so that replication can proceed without interference and that toxic recombination outcomes due to uncontrolled HJ resolution are minimized (Wild and Matos, 2016). Although deletion of BLM itself causes embryonic lethality in mice, individual loss of GEN1 or MUS81 does not confer a strong DNA repair-deficient phenotype in unperturbed cells, implying some degree of functional overlap between the two proteins (McDaniel et al., 2003; Sarbajna et al., 2014). Only when GEN1 and MUS81 are both absent is the genomic integrity of the cells severely subverted, resulting in compromised viability, gross chromosomal abnormalities, multinucleation, and heightened formation of micronuclei (Chan et al., 2018; Garner et al., 2013; Sarbajna et al., 2014; Wechsler et al., 2011).

Besides removing HR intermediates arising from recombinational DNA transactions, both SSEs process persistent replication intermediates to promote the completion of genome replication and sister chromatid disentanglement (Falquet and Rass, 2019). MUS81 (in complex with EME2) can cleave replication forks, structures resembling intact HJs such as four-way reversed forks, and D-loops to initiate and regulate replication fork restart via break-induced replication (BIR) (Hanada et al., 2007; Kikuchi et al., 2013; Mayle et al., 2015; Pepe and West, 2014a, 2014b). MUS81 is also essential for the ‘expression’ of common fragile sites (CFS)—sites that are prone to under-replication during stressed conditions—by cleaving stalled replication forks to enable POLD3-mediated mitotic DNA synthesis (MiDAS) (Debatisse et al., 2012; Minocherhomji et al., 2015; Naim et al., 2013; Ying et al., 2013). Generation of DSBs at stalled replication forks, however, is not a prerequisite for HR-dependent replication restart as the uncoupling of the DNA strands at collapsed forks generate ssDNA that can invade and re-initiate DNA synthesis (Lambert et al., 2010). Though GEN1 has been implicated in maintaining replication fork progression alongside MUS81 and in eliminating persistent replication intermediates in Dna2 helicase-defective yeast cells to potentially facilitate MiDAS, its importance in rectifying replication stress and DNA under-replication remains to be determined (Ölmezer et al., 2016; Sarbajna et al., 2014).

The role of GEN1 and MUS81 in replication-challenged *in vitro* settings employing ionizing irradiation, DNA-damaging agents, and replication inhibitors is well established. Less is known, however, about the importance of these HJ resolvases in an unperturbed *in vivo* context. Because B cells face an elevated risk of cell death and oncogenic transformation due to the high level of replication stress and DSBs inflicted on their genome by AID-dependent and -independent activities (Alt et al., 2013; Barlow et al., 2013; Basso and Dalla-Favera, 2015; Macheret and Halazonetis, 2015), we asked whether B cells require GEN1 and MUS81 to develop, survive, and perform their immunological functions. By employing different stage-specific Cre strains and a global *Gen1*-knockout mouse carrying floxed *Mus81* alleles, we report that the loss of *Gen1* and *Mus81* in early pro-B cells severely impaired B cell development whereas double-null mature B cells failed to form competent GCs at steady-state and after immunization. *Ex vivo* characterization of the double-knockout cells uncovered a proliferation block caused by G2/M arrest, potent activation of p53 and apoptotic pathways, induction of type I interferon (IFN) response, and widespread chromosomal aberrations. Our findings support the notion that the primary function of GEN1 and MUS81 in highly proliferative somatic cells such as B cells is in eliminating replication-derived HJ intermediates to ensure proper chromosome segregation and avert mitotic catastrophe.

## RESULTS

### GEN1 and MUS81 are critical to normal B cell lymphopoiesis

We analyzed a publicly accessible RNA-seq dataset (GSE72018) to ascertain the expression pattern of *Gen1* and *Mus81* in various developing and mature B cell subsets (Brazão et al., 2016). Expression of *Gen1* is higher in the proliferating pro-B, pre-B, and GC B cells than in follicular B, marginal zone B, and peritoneal B1a cells (**Figure 1A**). Conversely, *Mus81* RNA expression is relatively similar across all B cell subsets examined except in peritoneal B1a cells (**Figure 1B**). RT-qPCR analysis of primary splenic naïve B lymphocytes stimulated in culture with lipopolysaccharide (LPS) and interleukin-4 (IL-4) revealed a 10-fold increase in *Gen1* expression as early as 24 hours post-activation while the expression of *Mus81* was not altered following activation (**Figure 1–figure supplement 1A**). We surmise that *Gen1* expression is more closely associated with activation and proliferation of cells than is *Mus81* expression.

**Figure 1.**
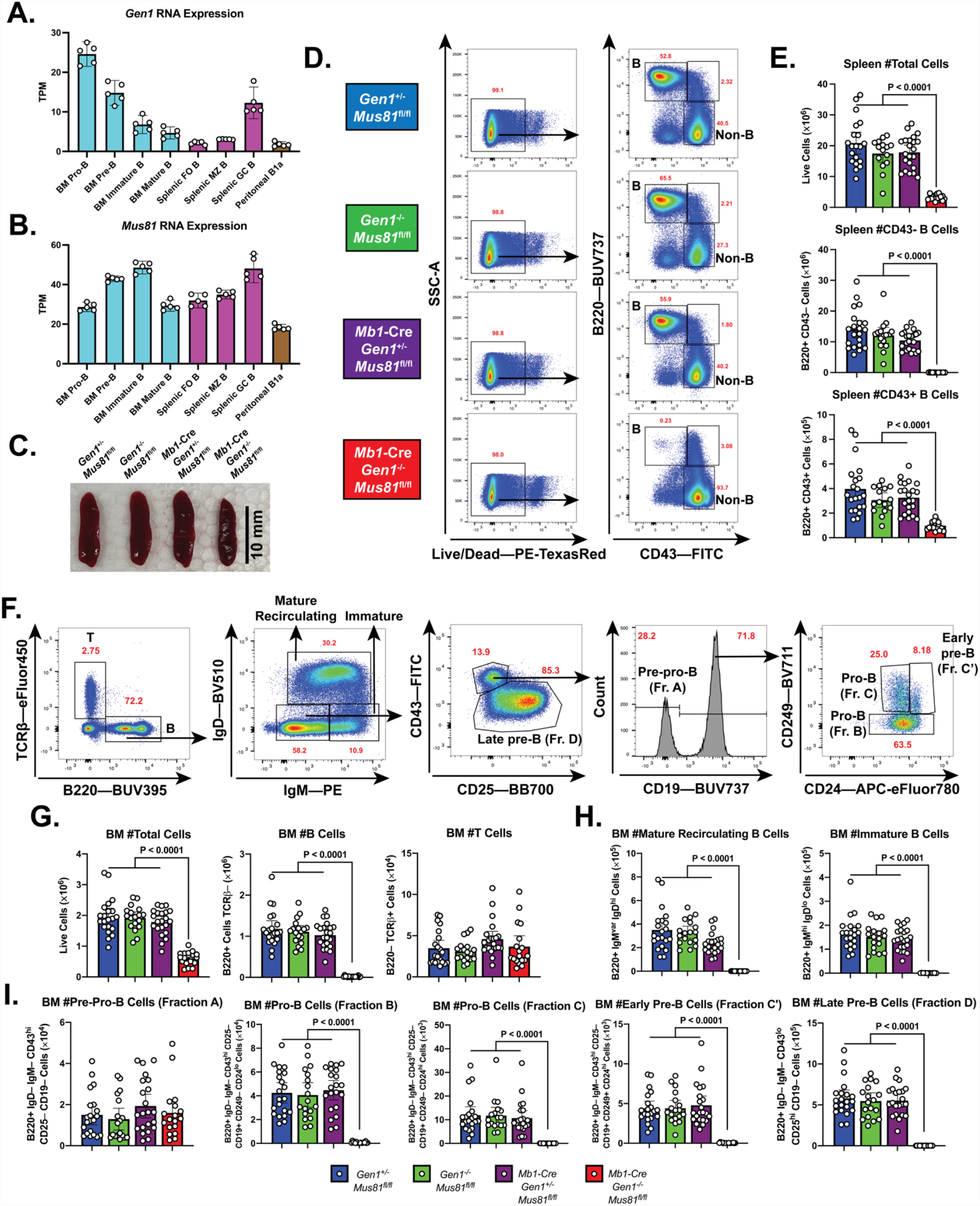
B cell development in the bone marrow and spleen of *Mb1*-Cre *Gen1*^-/-^ *Mus81*^fl/fl^ mice. (A and B) mRNA expression of *Gen1* (A) and *Mus81* (B) in developing and mature B cell subsets in the bone marrow (BM), spleen, and peritoneal cavity. FO: follicular, MZ: marginal zone (GEO Accession: GSE72018). (C) Spleens harvested from 4-month-old mice of the indicated genotypes. (D and E) Gating strategy (D) and absolute quantification (E) of live splenocytes, splenic B220+ CD43– B, and B220+ CD43+ B cells. (F) Gating strategy of total B (B220+ TCRβ–), T (B220– TCRβ+), mature recirculating, immature, pre-pro B (Fraction A), pro-B (Fractions B and C), early pre-B (Fraction C’), and late pre-B cells (Fraction D). (G–I) Absolute quantification of BM cellularity, total B, and total T cell populations (G), of mature recirculating and immature B cell populations (H), and of B cell populations belonging to fractions A to D (I). Data in (E) and (G)–(I) are from four independent experiments with 18 to 22 mice per genotype. Bars display the arithmetic mean and error bars represent the 95% confidence interval of the measured parameters. P-values were enumerated using ordinary one-way ANOVA analysis with Dunnett’s multiple comparisons test without pairing wherein all means were compared to the *Mb1*-Cre *Gen1*^-/-^ *Mus81*^fl/fl^ group. TPM, transcripts per million.

To interrogate the roles of GEN1 and MUS81 in B cell development and function, we generated *Gen1*^-/-^ mice by deleting the XPG nuclease domain encoded in exon 4 of *Gen1* and subsequently crossed them to *Mus81*^-/-^ mice (Dendouga et al., 2005) (**Figure 1–figure supplement 1B**). Whereas *Gen1*^-/-^ and *Mus81*^-/-^ mice were born in the expected Mendelian frequencies, no live *Gen1*^-/-^ *Mus81*^-/-^ pups were produced, indicating that constitutive loss of both GEN1 and MUS81 leads to embryonic lethality (**Figure 1–figure supplement 1C**). To circumvent this issue, we generated and bred mice harboring floxed *Mus81* alleles (*Mus81*^fl/fl^) to the *Mb1*-Cre strain, wherein Cre recombinase expression is driven by the *Cd79a* promoter and begins at the pre-pro B cell stage (Fahl et al., 2009; Hobeika et al., 2006). These mice were mated to *Gen1*^-/-^ *Mus81*^fl/fl^ mice to produce the conditional knockout *Mb1*-Cre: *Gen1*^-/-^ *Mus81*^fl/fl^, the single knockouts *Gen1*^-/-^ Mus81^fl/fl^ and *Mb1*-Cre: *Gen1*^+/-^ *Mus81*^fl/fl^, and the control *Gen1*^+/-^ *Mus81*^fl/fl^.

Analysis of the spleens of the *Mb1*-Cre *Gen1*^-/-^ *Mus81*^fl/fl^ mice found that the size of the total splenocyte population was reduced by ninefold compared with control and single knockout mice (**Figure 1C to E)**. The near complete ablation of mature B220+ CD43+ and B220+ CD43-cellular compartments indicated a significantly perturbed development or maintenance of peripheral B-lineage cells (**Figure 1E**). Examination of the bone marrow revealed a 75% reduction in cellularity that was caused by the severe loss of the total B cell population; the T cell numbers remained unaltered (**Figure 1F and G**). Quantification of the B-lineage subpopulations showed that both immature and mature recirculating B cells were absent (**Figure 1H**), and that the resolvase-deficient B cell progenitors produced few CD19^+^ CD43^+^ BP1^−^ CD24^var^ pro-B cells. (**Figure 1I**). The size of the pre-pro B cell population in these *Mb1*-Cre *Gen1*^-/-^ *Mus81*^fl/fl^ mice, conversely, was comparable to that of control and single-knockout mice, indicating that *Gen1* and *Mus81* in pre-pro B cells are necessary for differentiation into pro-B cells, the expansion and maintenance of pro-B cells, or both.

### Gen1 and Mus81 are required for robust germinal center responses

To circumvent the B cell developmental block in *Mb1*-Cre: *Gen1*^-/-^ *Mus81*^fl/fl^ mice, we bred the naïve B cell-specific deleter strain, *Cd23*-Cre, with *Gen1*^-/-^ *Mus81*^fl/fl^ mice to generate *Cd23*-Cre: *Gen1*^-/-^ *Mus81*^fl/fl^ (designated henceforth as DKO), *Cd23*-Cre: *Gen1*^+/-^ *Mus81*^fl/fl^ (*Mus81*-KO), *Gen1*^-/-^ *Mus81*^fl/fl^ (*Gen1*-KO), *Cd23*-Cre: *Gen1*^+/-^ *Mus81*^fl/+^ (Cre control), and *Gen1*^+/-^ *Mus81*^fl/fl^ (control) littermates. Though the B cell precursor subsets in the bone marrow of the DKO mice did not exhibit any major deficiencies in their frequencies or absolute numbers (**Figure 2–figure supplement 1A to E**), the total bone marrow cellularity was reduced by 30%, attributed to the 60% decrease in the number of the mature recirculating B cells (**Figure 2–figure supplement 1C**). In the DKO spleens, the frequencies and absolute numbers of the various B cell subsets were comparable to those of control and the *Gen1*-KO and *Mus81*-KO (collectively referred to as SKO) spleens (**Figure 2–figure supplement 1F to K**). RT-qPCR analysis of splenic mature DKO B cells activated with LPS+IL-4 confirmed the ablation of *Gen1* and *Mus81* transcript expression (**Figure 2–figure supplement 1L**). These data show that the deletion of *Gen1* and *Mus81* in the later stages of the B cell life cycle does not markedly impact the development and maintenance of homeostatic B cell compartments; thus, the DKO mice can serve as a genetic tool to investigate the functions of *Gen1* and *Mus81* in activated, mature B cells.

We next characterized the impact of *Gen1* and *Mus81* deletion on the steady-state GC response in the mesenteric lymph nodes and Peyer’s patches—sites where B cells continuously encounter and are activated by microbial and food antigens (**Figure 2A**). We found that the frequency and absolute number of GC B cells were decreased by 1.5 to 2-fold in DKO mice compared with control and SKO mice (**Figure 2B and C**). To assess the importance of GEN1 and MUS81 in supporting the formation and integrity of induced GCs, we immunized mice with sheep red blood cells (SRBCs) to elicit a T cell-dependent GC response. Mice were boosted 10 days after the initial dose and the GCs in the spleen were analyzed at day 14 (**Figure 2D**). Only in the DKO mice was GC formation abrogated: the size of the GCs was only 5% of that in control and SKO mice (**Figure 2E**). Such disruption in GC formation was also observed when DKO mice were challenged with another T-cell dependent antigen, NP-CGG (**Figure 2G**). Despite the compromised GC response, the frequencies and absolute numbers of total B220+ CD19+ B cells in both immunization settings were comparable in the DKO, SKO, and control littermates (**Figure 2F and H**). These experiments indicate that although *Gen1*-*Mus81*-null naïve B cells can persist in the periphery, they are unable to mount a productive GC reaction upon antigenic exposure at the barrier sites and in secondary lymphoid organs.

**Figure 2.**
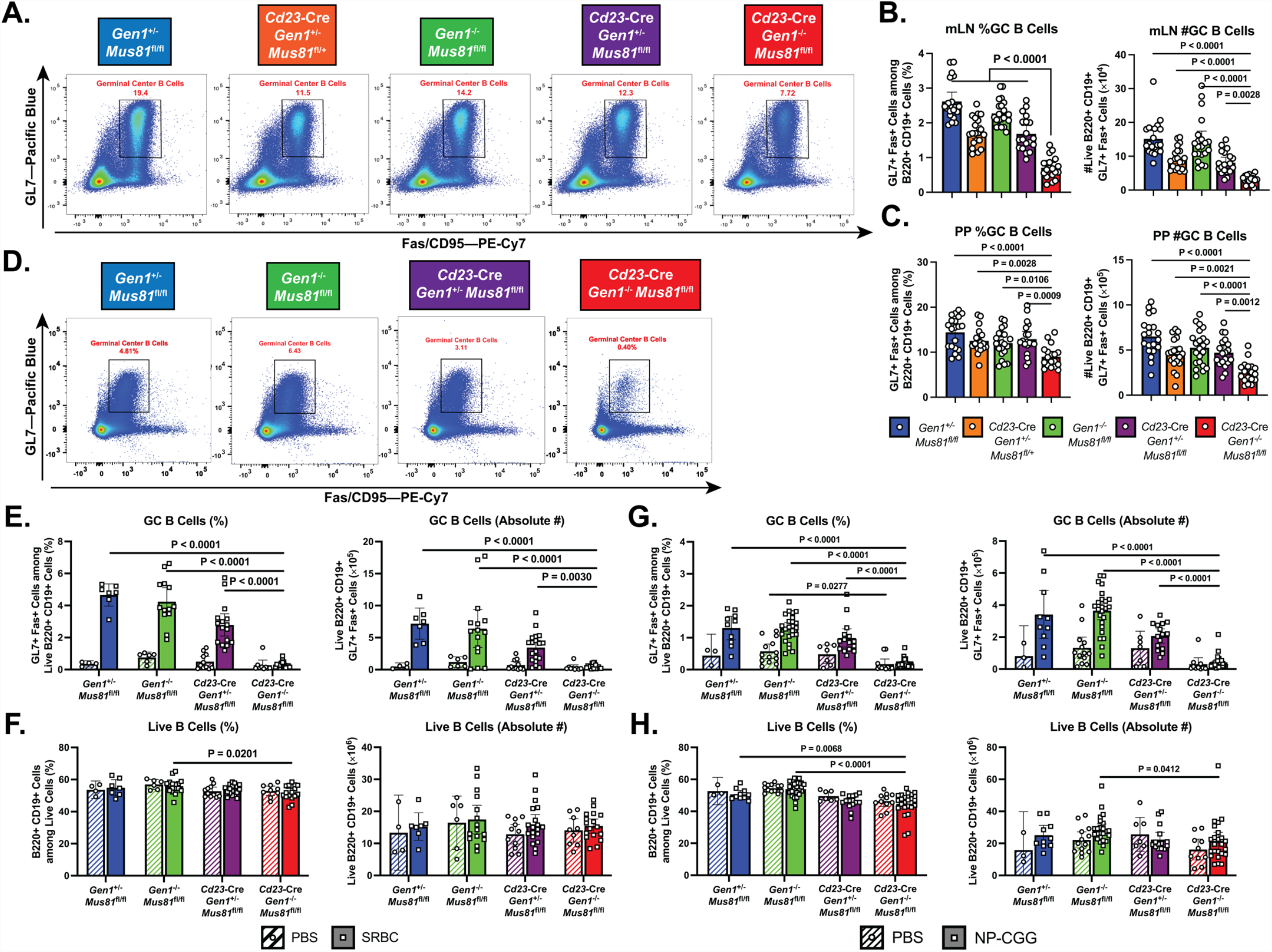
Homeostatic and induced GC responses in *Cd23*-Cre *Gen1*^-/-^ *Mus81*^fl/fl^mice. (A–C) Homeostatic GC response in mesenteric lymph nodes and Peyer’s patches. (A)Flow cytometric plots depicting the GL7+ Fas+ GC B cells in the Peyer’s patches of mice for each indicated genotype. (B and C) Quantification of the frequencies and absolute numbers of the GL7+ Fas+ GC B cell population in the mesenteric lymph node (B)and Peyer’s patches (C). (D–F) Evaluation of GC response during SRBC challenge. (D) Representative plots displaying the GL7+ Fas+ GC B cell population in the spleen of mice for each indicated genotype. (E and F) Quantification of the frequencies and absolute numbers of total B cells (E) and GC B cells (F) in the spleen of SRBC-immunized mice. (G and H) Assessment of induced GC response upon NP-CGG challenge in the *Cd23*-Cre *Gen1*^-/-^ *Mus81*^fl/fl^ mouse model at day 21 post-immunization. (G) Live cell frequencies and absolute numbers of live B220+ B cells in the spleen of PBS-treated and NP-CGG-immunized mice. (H) Quantification of the percentage and absolute count of GL7+ Fas+ GC B cells in the spleen. Data in (B) and (C) are from four independent experiments with 11 to 21 mice per genotype. Data in (D)–(F) are from three independent experiments with 4 to 9 mice per genotype for the PBS group and 7 to 20 mice per genotype for the SRBC group. Data in (G)–(H) are from three independent experiments with 5 to 13 per genotype in the PBS group and with 10 to 25 mice in the NP-CGG group. Bars represent the arithmetic mean and the error bars depict the 95% confidence interval of the measured parameters. For (A)–(C), P-values were computed by ordinary one-way ANOVA analysis with Dunnett’s multiple comparisons test without pairing in which the means were compared to the *Cd23*-Cre *Gen1*^-/-^ *Mus81*^fl/fl^ group. For (D)–(H), ordinary two-way ANOVA analysis with Dunnett’s multiple comparisons test without pairing was used to calculate the P-values. All means were compared within each treatment group to the *Cd23*-Cre *Gen1*^-/-^ *Mus81*^fl/fl^ cohort.

### GEN1 and MUS81 are necessary for B cell proliferation and survival

To mechanistically investigate the causes underlying the abrogated GC response in the DKO mice, we leveraged a tractable *ex vivo* culture system wherein purified splenic naïve B cells are induced to proliferate upon stimulation with various cocktails of mitogens and cytokines. The growth of splenic B cell cultures stimulated with LPS alone, LPS+IL-4 (LI), or LPS+TGF-β+anti-IgD dextran (LTD) was monitored by enumerating the live cells in culture using flow cytometry. Across all stimulation conditions, the DKO B cells were unable to expand—after 96 hours of culture, the number of live DKO cells was only 10% of that of control and SKO B cells (**Figure 3A and Figure 3–figure supplement 1A**). We then examined the proliferation dynamics of DKO B cells by labeling the cells with CellTrace™ Violet and tracking the dilution of the dye over time. We noted that at 72 hours post-LI and LPS simulations, between 10% and 15% of the live DKO cells remain undivided, indicating that a subset of DKO cells was incapable of proliferating (**Figure 3B and Figure 3–figure supplement 1B**). Moreover, only between 51% (LTD culture) and 68% (LPS culture) of the DKO cells that had proliferated underwent at least 3 rounds of cell division, in contrast to between 72% (LTD culture) and 86% (LPS culture) of the corresponding control and SKO populations (**Figure 3C and Figure 3–figure supplement 1B**).

**Figure 3.**
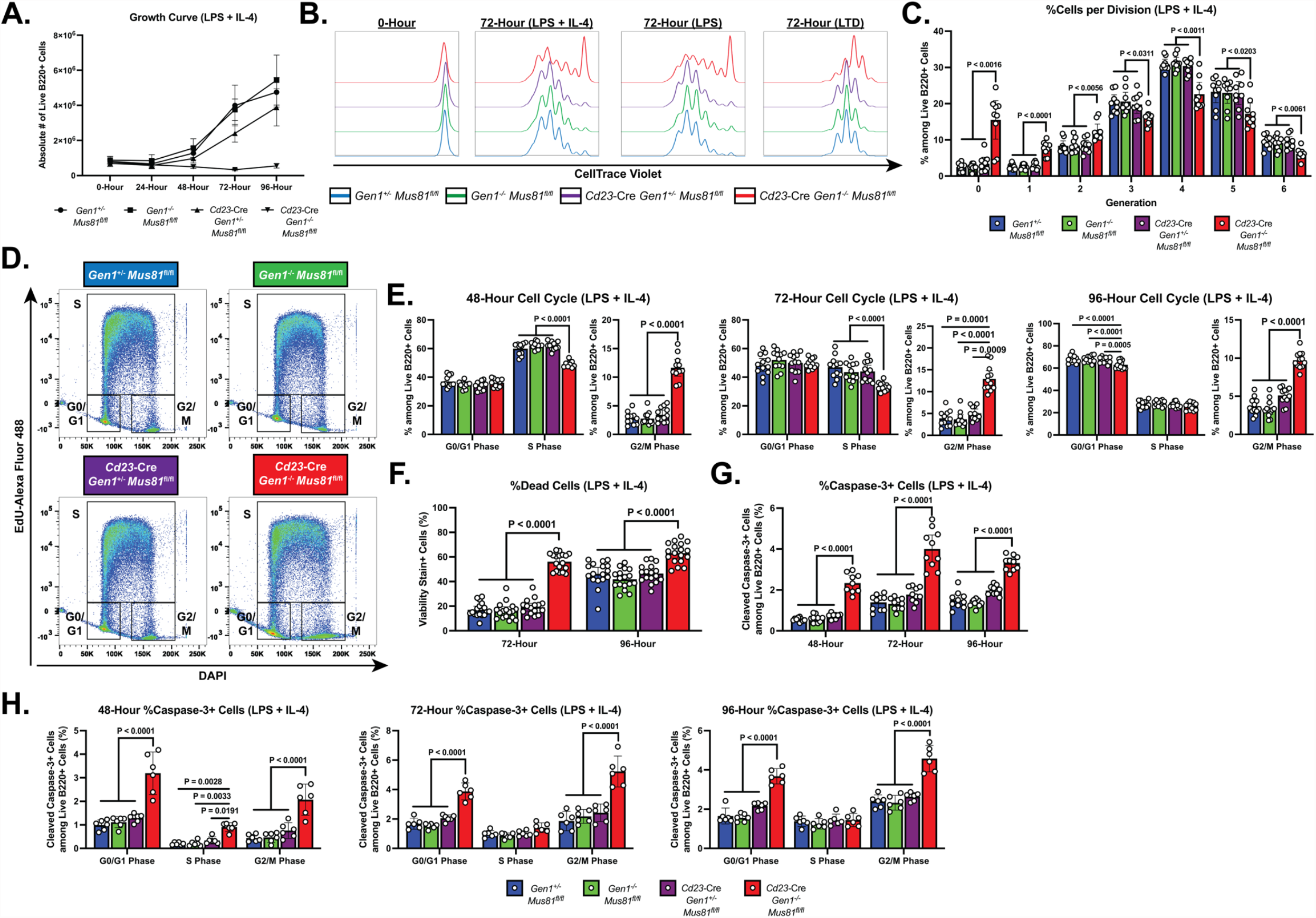
Growth, proliferation, cell cycle, and cell death profiles of *ex vivo*-stimulated DKO B lymphocytes. (A) Growth curve of LPS+IL-4 (LI)-activated B cell culture. (B) Representative CTV dilution profiles of *ex vivo*-activated B cells at 0 hour and 72 hours post-stimulation for all indicated genotypes and culture conditions. (C) Frequency of live B220+ cells in each division. (D) Representative flow cytometry plots delineating the cell cycle stages of the cultured B cells based on EdU positivity and nuclear DNA content as determined by the intensity of DAPI staining. (E) Frequency of live B cells in G0/G1, S, and G2/M phases. (F) Fraction of dead cells among B220+ singlets at 72 and 96 hours after LPS+IL-4 activation. (G) Percentage of caspase-3+ cells among live cells at 48, 72, and 96 hours post-stimulation. (H) Frequency of cleaved caspase-3+ cells among live B220+ cells in G0/G1, S, and G2/M phases after 48, 72, and 96 hours of culture. Data in (A) are from four independent experiments with 9 mice per genotype. Data in (B) and (C) are representative of four independent experiments with 7 to 9 mice per genotype. Data in (D) and (E) are from six independent experiments with 11 to 12 mice per genotype. Data in (F) are from seven experiments with 17 mice per genotype. Data in (G) are from five independent experiments with 9 to 10 mice per genotype. Data in (H) are from three independent experiments with 6 mice per genotype. Bars display the arithmetic mean and error bars represent the 95% confidence interval of the measured parameters. P-values were determined using ordinary two-way ANOVA analysis with Dunnett’s multiple comparisons test without pairing wherein the mean of the *Cd23*-Cre *Gen1*^-/-^ *Mus81*^fl/fl^ group was compared to the rest.

To gain further insight into the proliferation defect sustained by the activated DKO cells, we analyzed the cell cycle profile of these cells. Cultured B cells were pulse labeled with the nucleoside analog EdU to mark cells in S phase and stained with DAPI to quantify DNA content; the proportion of cells in G0/G1, S, and G2/M phases was determined by flow cytometry (**Figure 3D**). As early as 48 hours post-stimulation, the DKO cultures were enriched for cells in G2/M phase and depleted for cells in S phase (**Figure 3E**). This skewed cell cycle distribution of the DKO cells persisted up till 96 hours post-activation and was observed in all culture conditions (**Figure 3–figure supplement 1C and D**). Additionally, in the LTD culture, the fraction of DKO cells in the G0/G1 stage was reduced across all the time points examined (**Figure 3–figure supplement 1D**). Altogether, these results suggest that the dual loss of GEN1 and MUS81 causes the cells to stall in G2/M, impeding the completion of the cell cycle.

Aside from the proliferation deficiency, we also observed a 2 to 3-fold higher proportion of dead cells in the DKO cultures relative to that in control and SKO cultures (**Figure 3F and Figure 3–figure supplement 2A**). To ascertain whether elevated apoptosis contributed to the perturbed expansion of DKO B cell cultures, we quantified by flow cytometry the frequency of cells that stained for anti-cleaved caspase-3 antibody. As early as 48 hours post-stimulation, the proportion of caspase-3+ cells was 2- to 5-fold higher in DKO cultures compared with control and SKO cultures. The size of the caspase-3+ population peaked at 72 hours post-stimulation before declining (LI and LTD cultures) or remaining unchanged (LPS culture) by 96 hours post-stimulation (**Figure 3G and Figure 3–figure supplement 2B**). When we quantified the fraction of caspase-3+ cells in the different cell cycle phases, we found that across all the time points and stimulation conditions examined, DKO cells in G2/M and G0/G1 phases experienced a 2- to 5-fold higher level of apoptosis than their control and SKO counterparts (**Figure 3H and Figure 3–figure supplement 2C and D**). Taken together, these observations underscore an indispensable role for GEN1 and MUS81 in supporting the proliferative capacity and viability of activated B lymphocytes.

### Ablation of GEN1 and MUS81 induces p53 and type I interferon transcriptional programs

To assess the genome-wide transcriptional alterations underlying the proliferation and survival defects of *ex vivo*-activated DKO B cells, we conducted RNA-sequencing (RNA-seq) on activated B cells harvested at 48 hours post-stimulation, the time point at which the DKO cells were viable while displaying early signs of cell cycle perturbation and apoptosis. The RNA-seq analysis showed that the activated control and *Gen1*-KO B lymphocytes resemble each other transcriptomically, consistent with the lack of overt perturbations in *Gen1*-KO B cells (**Figure 4–figure supplement 1A**). Differential gene expression (DGE) analysis comparing *Mus81*-KO to control cells, however, identified 8 genes with a minimum of 2-fold upregulation in *Mus81*-KO cells (**Figure 4–figure supplement 1B**). The induction of only a few genes in *Mus81*-KO cells could be explained by the mild proliferation and survival perturbations that we had observed in our *in vivo* and *ex vivo* experiments. Between *Gen1*-KO and *Mus81*-KO cells, the transcript level of only 4 genes, including *Mus81* were differentially altered (**Figure 4–figure supplement 1C**). These findings collectively illustrate that the deletion of either *Gen1* or *Mus81* alone does not substantially alter the transcriptional landscape of activated B lymphocytes.

DGE analysis of control versus DKO cells, on the contrary, revealed that 279 genes were upregulated by at least twofold (log_2_ fold change ≥1; FDR <0.05) and 167 genes were downregulated by a minimum of log_2_ fold change of –0.3 in the DKO B cells (corresponding to a ≥19% de-enrichment relative to control cells) (**Figure 4A**). To identify the functional modules to which the differentially expressed genes in the DKO cells belong, we performed gene set enrichment analysis (GSEA) with the Hallmark Gene Sets from the Molecular Signatures Database (MSigDB) (Liberzon et al., 2015). We identified the p53 pathway (25/200 genes with log_2_ fold change ≥1; e.g., *Ccng1, Zfp365, Plk2, Phlda3, Cdkn1a, Zmat3*) and apoptosis (15/161 genes with log_2_ fold change ≥0.5) signatures among the top 5 enriched gene sets whereas gene sets containing genes targeted by *Myc* and E2F transcription factors and genes involved in progression through the G2/M checkpoint were among the most de-enriched in the DKO cells (**Figure 4B and C and Figure 4–figure supplement 1D to F**), concurring with our findings that DKO cultures exhibit perturbed cell cycle progression, G2/M arrest, proliferation irregularities, and heightened apoptosis. We also noted that the *ex vivo*-activated DKO B cells displayed a robust type I IFN gene signature, as exemplified by the high enrichment score (NES = 2.35; FDR = 6.34 × 10^−3^) and the induction (log_2_ fold change ≥0.5) of 21/97 genes in the gene set (e.g., *Ifit3, Ifit3b, Ifitm3, Mx1, Rsad2*) (**Figure 4C**). We posit from this analysis that the deficiency of both GEN1 and MUS81 in cells activates p53-dependent pathways to arrest cell cycle for the repair of genomic insults sustained during cellular growth and proliferation, and to initiate apoptosis when such DNA lesions are beyond tolerance and repair.

**Figure 4.**
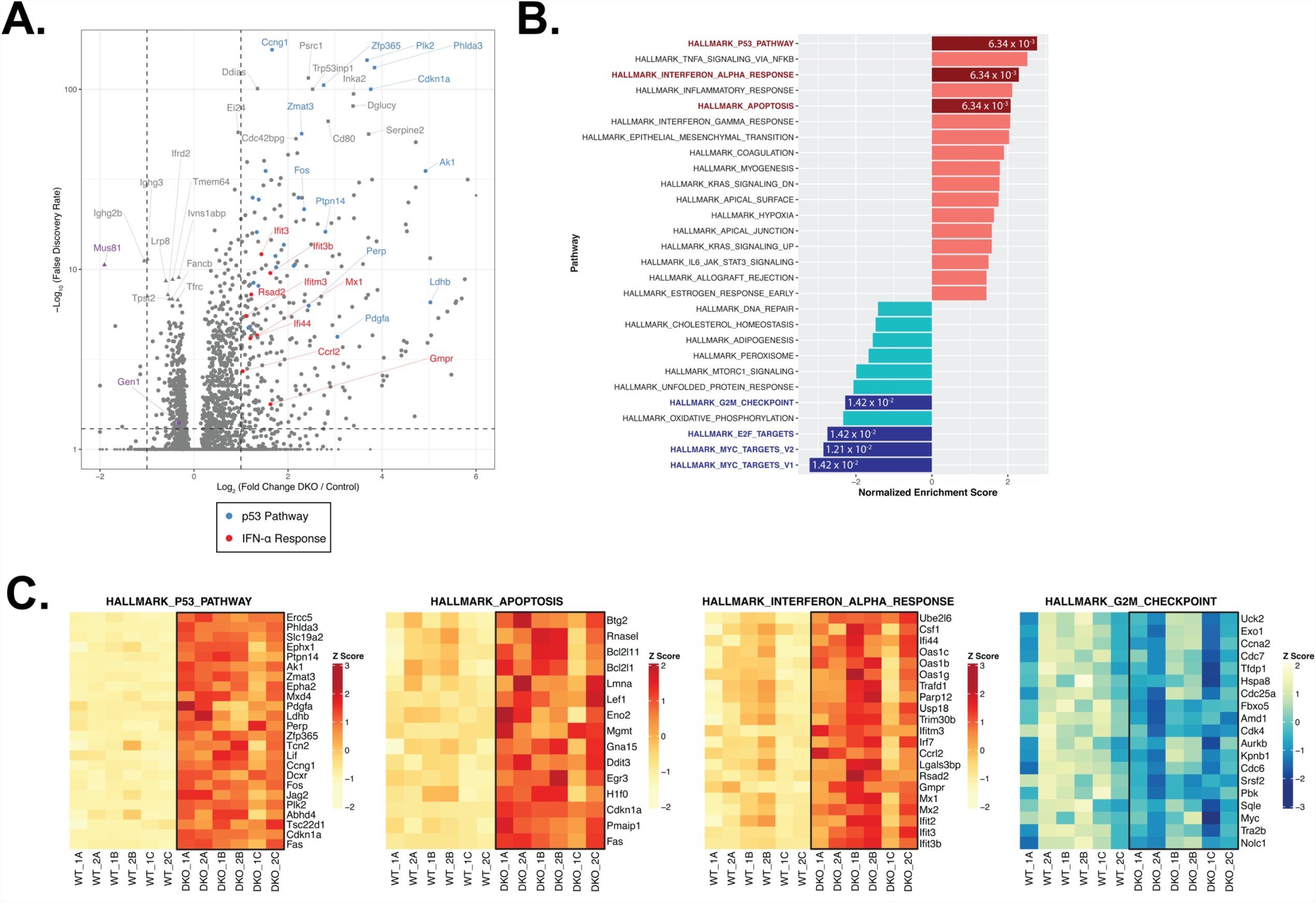
RNA-seq and GSEA analyses of activated control and DKO B cells. (A) Volcano plot depicting the differential gene expression between control and DKO cells. Labeled squares indicate the top 15 most significantly upregulated genes and the labeled triangles are the 10 most downregulated genes in DKO cells. Using the Hallmark Gene Sets as a reference, genes labeled in blue are categorized as genes in the p53 pathway while those labeled in red are defined as interferon alpha (IFN-α) response genes. Purple triangles mark *Gen1* and *Mus81*. (B) Graph depicting the list of Hallmark Gene Sets that are differentially expressed between control and DKO cells (FDR < 0.05). Value in each bar denotes the FDR for that gene set. (C) Heatmaps displaying the relative expression of genes within the indicated Hallmark Gene Sets that meet the expression cutoff (log2 fold change >1 for p53 pathway; >0.5 for apoptosis and interferon alpha response; <–0.3 for G2/M checkpoint; with FDR < 0.05). Data are from 6 mice per genotype. The labels ‘1’ and ‘2’ represent male and female mice, respectively, and ‘A’ to ‘C’ indicate the three experimental *ex vivo* groups.

### GEN1 and MUS81 maintain the genome stability of activated B lymphocytes

GEN1 and MUS81 resolve structural intermediates of HR to prevent genome destabilization caused by the toxic accumulation of erroneously processed recombination structures (Chan et al., 2018; Sarbajna et al., 2014). Reasoning that the genomic integrity of proliferating DKO B cells might similarly be compromised, we prepared metaphase spreads from *ex vivo*-stimulated B cells for evaluation of chromosomal integrity. DKO B cells displayed approximately 6 times as many abnormalities per metaphase as control and SKOs B cells (**Figure 5A and B**). Furthermore, whereas almost 95% of control and SKO metaphases had no more than 2 aberrations, 45% of DKO metaphases showed 3 or more chromosomal defects (**Figure 5C**). All forms of aberrations including breaks, DNA fragments, fusions, and radials were detected at an elevated rate in DKO B cells compared with control and SKOs cells (**Figure 5D**). Notably, almost 70% of the breaks observed were chromosome-type breaks that occur at the same position on both sister chromatids (**Figure 5E**). To better understand the nature and origins of the chromosomal irregularities occurring in the activated DKO B cells, we performed telomere fluorescence in situ hybridization (Tel-FISH). In DKO cells, the breakage occurred proximal to the telomeres, resulting in paired DNA fragments containing telomeric DNA (**Figure 5F**). Such symmetrical breakage has been proposed to arise from unresolved recombination intermediates (Garner et al., 2013). These data argue that GEN1 and MUS81 eliminate complex recombination intermediates in a timely fashion to preserve the chromosomal integrity of proliferating B lymphocytes.

**Figure 5.**
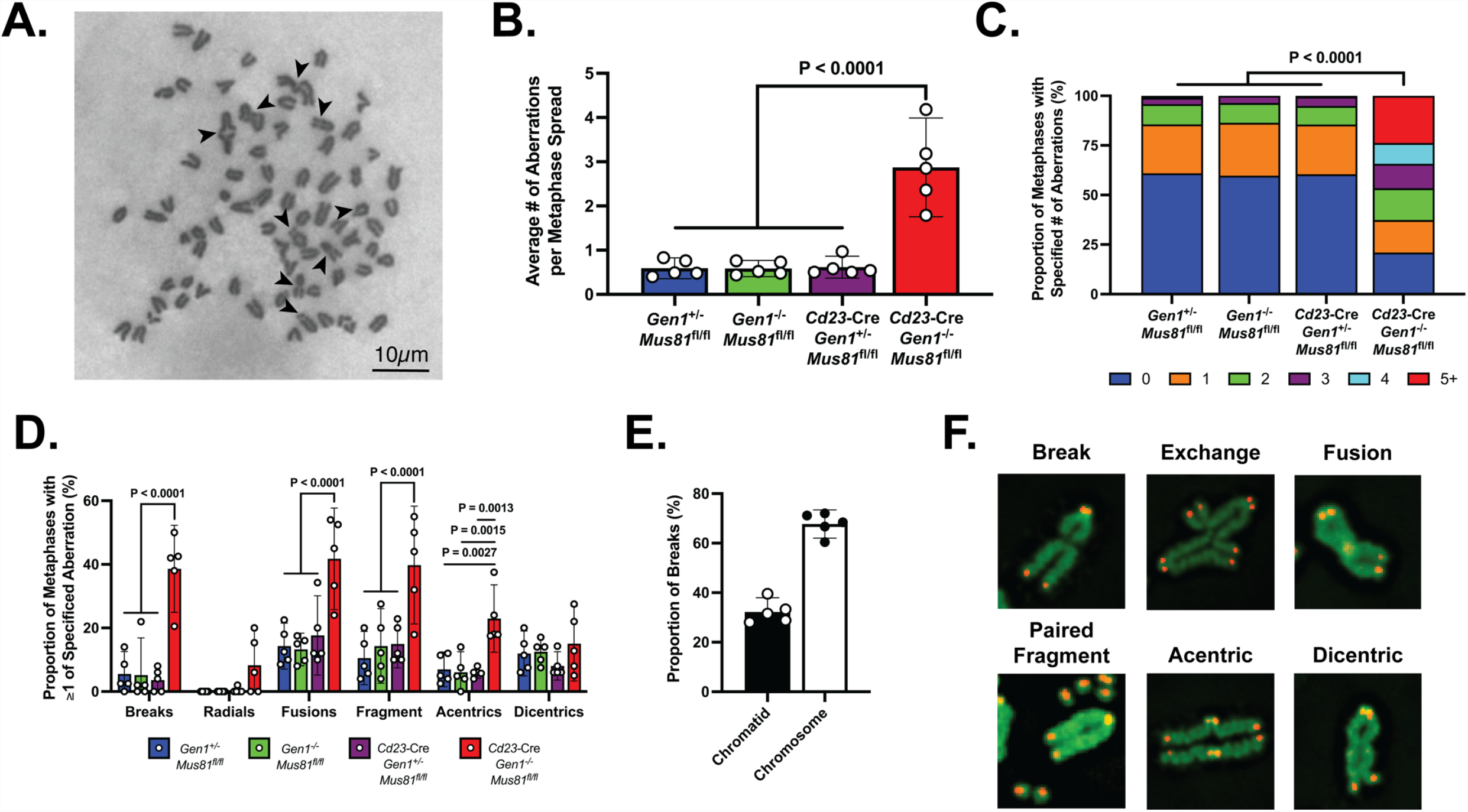
Metaphase chromosomal analysis of activated DKO B cells. (A) Representative image of a DKO metaphase spread with arrows indicating chromosomal breaks, fragments, and fusions in metaphases of activated DKO B cells. (B) Quantification of the average number of chromosomal aberrations across 45 to 50 metaphase spreads prepared from each B cell culture. (C) Percentage breakdown of metaphases exhibiting 0 to greater than 5 chromosomal aberrations. (D) Fraction of metaphases containing the different types of chromosomal abnormalities. Total percentage per genotype exceeds 100% as some metaphases exhibit more than one type of abnormality. (E) Proportion of chromatid and chromosome breaks among the 163 breaks observed in DKO metaphase spreads exhibiting at least one break. (F) Immunofluorescence images of Tel-FISH of metaphases of DKO B cells highlighting the proximal location of the chromosomal damage to the telomeres. Data in (A–D) are from three independent experiments with 5 mice (totaling between 215 and 235 metaphase spreads) per genotype. For (C) and (D), the percentage values are the average of the data combined from all 5 mice. Bars display the arithmetic mean and error bars represent the 95% confidence interval of the measured parameters. P-values were computed with ordinary one-way ANOVA analysis (B and D) and with the Kruskal-Wallis test (C) with Dunnett’s multiple comparisons test without pairing. Means of all groups were compared to that of *Cd23*-Cre *Gen1*^-/-^ *Mus81*^fl/fl^.

## DISCUSSION

HR constitutes one of the two major pathways cells utilize to repair DSBs. This mode of repair is largely restricted to late S and G2 phases as it requires the sister chromatid as a template to restore the fidelity of the damaged DNA strand (Heyer, 2015). HJ intermediates generated in this process are dissolved by the BTR complex and resolved by GEN1 and the SLX1-SLX4-MUS81-EME1 complex (West et al., 2016). Failure to eliminate HJs prohibits chromosomal segregation during mitosis, interfering with the faithful transmission of genetic material to daughter cells while yielding aberrant mitotic structures including bulky chromatin bridges and ultrafine anaphase bridges (UFBs) that threaten chromosomal integrity and genomic stability (Chan and West, 2018; Sarlós et al., 2017; West et al., 2016). B cells encounter a diverse array of genotoxic stresses throughout their life cycle, of which DSBs—both spontaneously and deliberately generated—are considered among the most deleterious lesions to the cells’ genome. As such, B cells serve as an informative physiological platform to interrogate the DNA transactions influencing genome integrity.

We established a *Gen1*^-/-^ mouse model possessing *Mus81*^fl/fl^ alleles to examine the roles of the HJ resolvases GEN1 and MUS81 at various stages of B cell development. Inactivation of both resolvases in early B cell precursors abolished the production of mature B cells in the bone marrow and periphery, whereas ablation in naïve mature B cells impaired germinal center formation. *Ex vivo* cellular and transcriptomics analyses reveal that these resolvase-deficient B cells exhibited significant proliferation and viability perturbations underpinned by widespread chromosomal abnormalities and activation of p53-dependent cell cycle arrest and apoptosis.

The severe attrition of pro-B cells in the *Mb1*-Cre *Gen1*^-/-^ *Mus81*^fl/fl^ mice could be caused by a developmental arrest during the transition from pre-pro B cell to pro-B cell stage, by proliferation and survival defects of the pro-B cells, or both. Pro-B cells, compared to the mitotically inactive pre-pro B cells, undergo IL-7R-dependent expansion prior to V(D)J recombination and so are more likely to suffer from intense replication stress that could necessitate commensurate effort of replication restart (Hardy et al., 1991; Peschon et al., 1994). As resolvase-knockout pro-B cells are unable to resolve HJ intermediates that arise from such restart activities, they would fail to thrive and proliferate further. Similarly, the lack of robust chronic and induced GC responses in the *Cd23*-Cre *Gen1*^-/-^ *Mus81*^fl/fl^ mice can be ascribed to the inability of activated DKO cells to undergo sustained proliferation due to the combination of a profound block at the G2/M transition and p53-activated apoptosis, as shown by our cell cycle and RNA-seq analyses. DKO cells not only have a lower propensity to initiate proliferative bursts, but are also incapable of executing as many divisions as resolvase-sufficient B cells when activated. We did not observe a time-dependent G2/M accumulation in the DKO culture, suggesting that the arrested DKO cells could not tolerate for long periods the high level of DNA damage sustained and consequently undergo mitotic catastrophe, evidenced in the comparatively higher rate of apoptosis among G2/M cells. The inability of the BTR complex to compensate for the absence of GEN1 and MUS81 and maintain the viability of proliferating DKO B cells thus suggests that DNA replication generates persistent double HJs and multiple BTR-refractory HJ species such as single HJs and nicked HJs that must be processed and eliminated by these resolvases to suppress genomic instability and catastrophic mitosis (García-Luis and Machín, 2014). Resting mature DKO cells, however, persisted unscathed in the spleen, as exemplified by the normal frequencies and numbers of splenic B cell subpopulations in the *Cd23*-Cre *Gen1*^-/-^ *Mus81*^fl/fl^ mice. Splenic naïve B cells do not self-renew; instead, they are constantly replenished from immature precursors produced in the bone marrow (Hao and Rajewsky, 2001). Hence, resting mature B cells are spared from replication-derived genotoxicity, rendering GEN1 and MUS81 dispensable in these cells.

Our observation of extensive chromosomal aberrations in the genetic resolvase-null mouse B cells concur with that in Sarbajna et al. (2014) employing siRNA depletion of *Gen1* and *Mus81* in cells treated with replication inhibitors. Recombination intermediates generated in S phase that fail to be resolved in the absence of GEN1 and MUS81 evade the checkpoint response and persist into mitosis, generating homologous recombination ultrafine bridges (HR-UFBs) (Chan et al., 2018; Mohebi et al., 2015; Tiwari et al., 2018). Breakage of the HR-UFBs during cytokinesis activates the DNA damage checkpoint in the next cell cycle, triggering non-homologous end joining-mediated fusion of DNA ends that leads to widespread chromosomal rearrangements (Chan et al., 2018). The preponderance of chromosome breaks that appear to occur at identical sites on sister chromatids suggests that the aberrations result from defective resolution of inter-chromatid recombination intermediates (Kikuchi et al., 2013; Shimizu et al., 2020; Wechsler et al., 2011). This phenotype is distinct from that of irradiated HR-defective mutants in which chromatid breaks predominate (Fujita et al., 2013; Shimizu et al., 2020). We cannot completely exclude the role of MUS81—and potentially GEN1—in initiating BIR and MiDAS at CFS and other under-replicated regions where their absence leads to the formation of FANCD2-flanking fragile site-UFBs (FS-UFBs) between the segregating chromatids (Naim et al., 2013; Ying et al., 2013). However, the frequent occurrence of breaks at corresponding locations on paired sister chromatids argues for a failure of DKO B cells to resolve HR intermediates. Visualization of UFBs in these cells may be helpful in defining the molecular origins of these unique aberrations. Detailed investigations to ascertain whether HR-UFBs derive predominantly from genomic loci where stalled replication forks are preferentially restarted through recombination-dependent mechanisms could provide mechanistic insights into the genomic instability of resolvase-null B cells and clarify the manner by which GEN1 and MUS81 resolve replication stress— do they cleave persistent and late-occurring replication intermediates such as reversed forks or do they process HR intermediates (e.g., HJs) arising from recombination-mediated restart of perturbed forks? Further studies to determine whether the loci where HR-UFBs manifest encompass early replication fragile sites (ERFS) should be pursued given that such sites have recently been identified as hotspots for transcription-replication conflicts and breakpoints of chromosomal rearrangements in B lymphocytes (Barlow et al., 2013).

We posit that the persistence of unresolved HR intermediates generated following recombination-mediated fork restart leads to the manifestation of aberrant mitotic structures including chromatin bridges, HR-UFBs, and micronuclei that enable rampant fusions of broken chromosomes and amplification of replication-associated DNA damage during the next cell cycle. Our studies show that such extensive chromosomal instability and genomic damage consequently activate p53-dependent G2/M arrest and apoptosis. Concurrently, these resolvase-null B cells exhibit a type I IFN transcriptional signature, potentially a ramification of high cytoplasmic levels of self-DNA. The synthetic lethality of GEN1 and MUS81 deficiencies in B cells highlights the essentiality of structure-selective endonucleases in eliminating replication-derived recombination intermediates to safeguard the genomic integrity of proliferating cells and to permit the sustained proliferation and survival capacities required for the proper development and functionality of B lymphocytes, and possibly other immune cells.

## ACKNOWLEDGEMENTS

We would like to thank the past and present members of the Chaudhuri and the Petrini labs for technical assistance, productive discussions, and constructive feedback. J.C. was supported by grants from the NIH (R01AI072194, R01AI124186, R56AI072194, U54CA137788, and P30CA008748), the Starr Cancer Research Foundation, the Ludwig Center for Cancer Immunotherapy, MSKCC Functional Genomics, and the Geoffrey Beene Cancer Center. J.H.J.P. was supported by grants from the NIH (R01GM56888, R35GM136278, U54OD020355, and P30CA008748). We thank A. Bravo for help with maintenance of the mouse colony. We acknowledge the use of the Memorial Sloan Kettering Cancer Mouse Genetics Core Facility.

## AUTHOR CONTRIBUTIONS

Keith Conrad Fernandez, Conceptualization, Methodology, Investigation, Formal Analysis, Visualization, Writing – Original Draft Preparation, Writing – Review & Editing; Laura Feeney, Methodology, Investigation, Formal Analysis, Visualization, Writing – Original Draft Preparation; Ryan M. Smolkin, Software, Formal Analysis, Writing – Review & Editing; Allysia J. Matthews, Methodology, Investigation, Formal Analysis; Wei-Feng Yen, Methodology, Investigation, Formal Analysis; John H.J. Petrini, Conceptualization, Resources, Writing – Review & Editing, Supervision, Project Administration, Funding Acquisition; Jayanta Chaudhuri, Conceptualization, Resources, Writing – Review & Editing, Supervision, Project Administration, Funding Acquisition

## MATERIALS AND METHODS

### Mice

*Gen1*^-/-^ mice were generated at the Memorial Sloan Kettering Cancer Center Mouse Genetics Core Facility. *Mus81*^fl/fl^ mice were generated using ES cell clone purchased from EUCOMM. Experiments were performed using mice between 8- and 16-week-old. When littermate controls are not available, age-matched controls were employed in experiments. All mice were housed and maintained in groups of five under specific pathogen-free conditions, and euthanized at the time of analyses in accordance with guidelines for animal care established by Memorial Sloan Kettering Cancer Center Research Animal Resource Center and the Institutional Animal Care and Use Committee (IACUC).

### Flow cytometry

Single-cell suspensions were prepared from mouse spleen, mesenteric lymph nodes, and Peyer’s patches by pressing through a 70 μm cell strainer (Corning), and bone marrow cells were harvested from the tibia. Splenic and bone marrow suspensions were resuspended in red blood cell lysis buffer (150 mM NH_4_Cl, 10 mM KHCO_3_, 0.1 mM EDTA) for 5 minutes at room temperature and then neutralized with B cell media (RPMI 1640 with L-glutamine (Gibco) supplemented with 15% fetal bovine serum (Corning), 1% penicillin-streptomycin (GeminiBio), 2 mM L-glutamine (Memorial Sloan Kettering Cancer Center Media Preparation Facility), and 55 μM β-Mercaptoethanol (Gibco)). After washing with PBS, cells were stained with Zombie Red™ fixable viability dye (BioLegend) and rat anti-mouse CD16/CD32 Fc Block (BD Biosciences), followed by staining with antibodies for cell surface markers at 4°C for 30 minutes. The following antibodies and their respective clones were used in this study: B220 (RA3-6B2), CD19 (ID3), TCRβ (H57-597), CD43 (R2/60), IgD (11.26.2a), IgM (II/41; polyclonal), CD25 (PC61), CD249 (BP-1), c-Kit (2B8), CD93 (AA4.1), CD24 (M1/69), CD138 (281-2), CD21/CD35 (7E9), CD23 (B3B4), GL7 (GL7), CD95 (Jo2), and CD38 (90). All antibodies were purchased from BD Biosciences, eBioscience, and BioLegend. For intracellular cleaved caspase-3 staining, *ex vivo*-stimulated cells were stained with cell surface markers followed by staining with anti-cleaved caspase-3 antibody (C92-605; BD Biosciences) for 45 minutes at 4°C after processing with Fixation/Permeabilization kit (BD Biosciences) according to the manufacturer’s protocol. Data was obtained using an LSR II flow cytometer (BD Biosciences) and analyzed with FlowJo 10.6 (BD Biosciences).

### Immunization

For SRBC immunization, packed SRBCs (Innovative Research) were washed thrice with PBS, counted with hemocytometer, and resuspended to a concentration of 10 million cells/μL. 500 million cells were then administered intraperitoneally. Mice were boosted with the same number of SRBCs on day 10 before spleens were harvested for analysis on day 14. For NP-CGG immunization, mice were injected intraperitoneally with 100 μg NP-CGG (ratio 30–39; Biosearch Technologies) resuspended in Imject™ Alum adjuvant (Thermo Scientific). Mice were boosted on day 14 and euthanized on day 21 for analysis of immune response in the spleen.

### Primary B cell *ex vivo* stimulation

Splenic B cells were harvested and processed into single-cell suspensions by pressing through a 70 μm cell strainer. Naïve B cells were then purified by negative selection using anti-CD43 microbeads (Miltenyi Biotec) according to the manufacturer’s protocol. B cells were plated at a density of 1 × 10^6^ cells/mL in B cell media in a 6-well dish. B cells were then stimulated with one of the following cytokine cocktails: 30 μg/mL LPS (Sigma); 30 μg/mL LPS plus 25 ng/mL IL-4 (R&D Systems); or 5 μg/mL LPS, 2 ng/mL recombinant human TGF-β1 (R&D Systems), and 333 ng/mL anti-IgD dextran conjugates (Fina Biosolutions). Cultures were split by half at 48-hour and 72-hour post-stimulation.

### qPCR analysis

Total RNA was harvested from 48-hour *ex vivo* B cell cultures using Quick-RNA™ Microprep Kit (ZymoResearch) and reverse transcribed to cDNA using High-Capacity cDNA Reverse Transcription Kit (Applied Biosystems). TaqMan™ probes specific for *Gen1* (Mm00724023_m1), *Mus81* (Mm00724023_m1), and *Ubc* (Mm01201237_m1) were used to amplify the cDNA transcripts. qPCR experiments were performed with the TaqMan™ Fast Advanced Master Mix (Applied Biosystems) in a 384-well format using an Applied Biosystems QuantStudio 6 Flex instrument. Relative gene expression was calculated using the 2^-ΔΔCT^ method and normalized to *Ubc* expression.

### Proliferation analysis

Purified naïve splenic B cells were stained with 5 μM CellTrace™ Violet (Invitrogen) in PBS for 20 minutes at room temperature in the dark. Cells were washed with B cell media to quench the dye before resuspension in fresh B cell media and subsequent incubation for at least 10 minutes at 37°C. Equal labeling between the genotypes was verified by flow cytometry immediately after labeling. Cytokine cocktails were then added to the B cell cultures to initiate stimulation.

### Cell cycle analysis

Prior to flow cytometric analysis, *ex vivo* B cells were treated with 10 μM EdU for 1 hour. Cells were harvested and washed with PBS before staining with antibodies for surface proteins. Cells were then processed using Click-iT™ EdU Alexa Fluor™ 488 Flow Cytometry Assay Kit (Invitrogen) according to the manufacturer’s protocol. Cells were subsequently stained with FxCycle™ Violet Stain (Invitrogen) for 15 minutes at room temperature in the dark before flow cytometry.

### RNA-sequencing library generation and analyses

B cells were cultured for 48 hours before total RNA was extracted using Quick-RNA™ Microprep Kit (ZymoResearch) and mRNA was isolated using the NEBNext® Poly(A) mRNA Magnetic Isolation (New England BioLabs). Stranded Illumina libraries were prepared with Swift Rapid RNA Library Kit according to the manufacturer’s instructions (Swift Biosciences). Indexed libraries were sequenced on a HiSeq X Ten platform, and an average of 30 million 150-bp paired-end reads were generated for each sample (Novogene, Beijing, China). The resulting FastQ files were processed to remove adapters and low-quality reads, using GATK v4.1.9.0 (Auwera et al., 2013). STAR v2.7.7a (Dobin et al., 2013) aligned the reads to GRCm38.p6 and gencode vM25 (Frankish et al., 2021), and GATK removed the duplicates. A count matrix was generated using featureCounts v2.0.1 (Liao et al., 2014), and DESeq2 v1.30.1 (Love et al., 2014) generated differential expression matrices. ggPlot2 v3.3.4 (Wickham, 2016) was used for volcano plots, highlighting genes that fall in the designated areas (see text). fgsea v1.16.0 (Korotkevich et al., 2021) and msigdb h.all.v7.4 (Liberzon et al., 2015; Subramanian et al., 2005) analyzed the differentially expressed genes (those with FDR <0.05), in DKO cells compared to control cells to determine which gene sets were enriched and de-enriched. Then, ggplot2 and a modified fgsea script was used to generate GSEA plots. Lastly, the feature count matrix was also used to produce normalized TPM values for all genes in each sample; these were then plotted with ComplexHeatmap v2.6.2 (Gu et al., 2016). All scripts are deposited in GitHub (Smolkin R., 2022). For the analysis of GSE720181, the dataset was downloaded as a featureCounts matrix and converted to TPM values in R v4.0.5.

### Metaphase spreads

Metaphase chromosome spreads were prepared by incubating cells with 100 μg/mL KaryoMAX™ Colcemid™ Solution in PBS (Gibco) for 3 hours. Cells were harvested at 1000 rpm and resuspended in 75 mM KCl at 37°C for 15 minutes. Cells were fixed in a 3:1 mixture of ice-cold methanol/acetic acid at least overnight at –20°C. Samples were then dropped onto pre-cleaned slides, briefly steamed (<5 seconds) over an 80°C water bath to disperse nuclei and air-dried overnight at room temperature. Slides were Giemsa stained and mounted using Fisher Chemical™ Permount™ Mounting Medium (Fisher Scientific). Images were acquired on an Olympus IX50-S8F microscope using a 100x objective and images were analyzed using ImageJ.

### Telomere FISH

Metaphase chromosome spreads were prepared and dropped onto slides as described above. Instead of Giemsa staining, samples were treated with 100 μg/mL RNAse A for 1 hour at 37°C, dehydrated with a series of 70%, 90% and 100% ethanol for 5 minutes each at room temperature, then allowed to air dry. Hybridization with 0.5 μg/mL CY-3 (CCCTAA)_3_ probe (PNA Bio) was carried out in hybridization buffer (10mM Tris pH 7.5, 70% formamide, 0.5% blocking reagent (Roche)). Samples were denatured at 75°C for 5 minutes then hybridization was allowed to proceed at room temperature for 16 hours. Slides were washed twice in wash buffer (10 mM Tris pH 7.5, 0.1% BSA, 70% formamide) and then three times in PBS/0.15% Triton X. Slides were then incubated for 10 minutes at room temperature in SYTOX™ Green Nucleic Acid Stain (diluted to 0.5 mM in PBS). After a final PBS wash, slides were mounted with ProLong™ Gold Antifade Mountant with DAPI (Invitrogen). Images were acquired on a DeltaVision Elite Cell Imaging System (GE Healthcare Life Sciences) with a CMOS Camera on an Olympus IX-71 microscope using a 60X objective. Images were analyzed using ImageJ.

### Statistical analysis

Graphical representation of data and statistical analyses were performed using Prism 9 (GraphPad Software). Tables were prepared using Numbers 11 (Apple).

**Figure 1–figure supplement 1.**
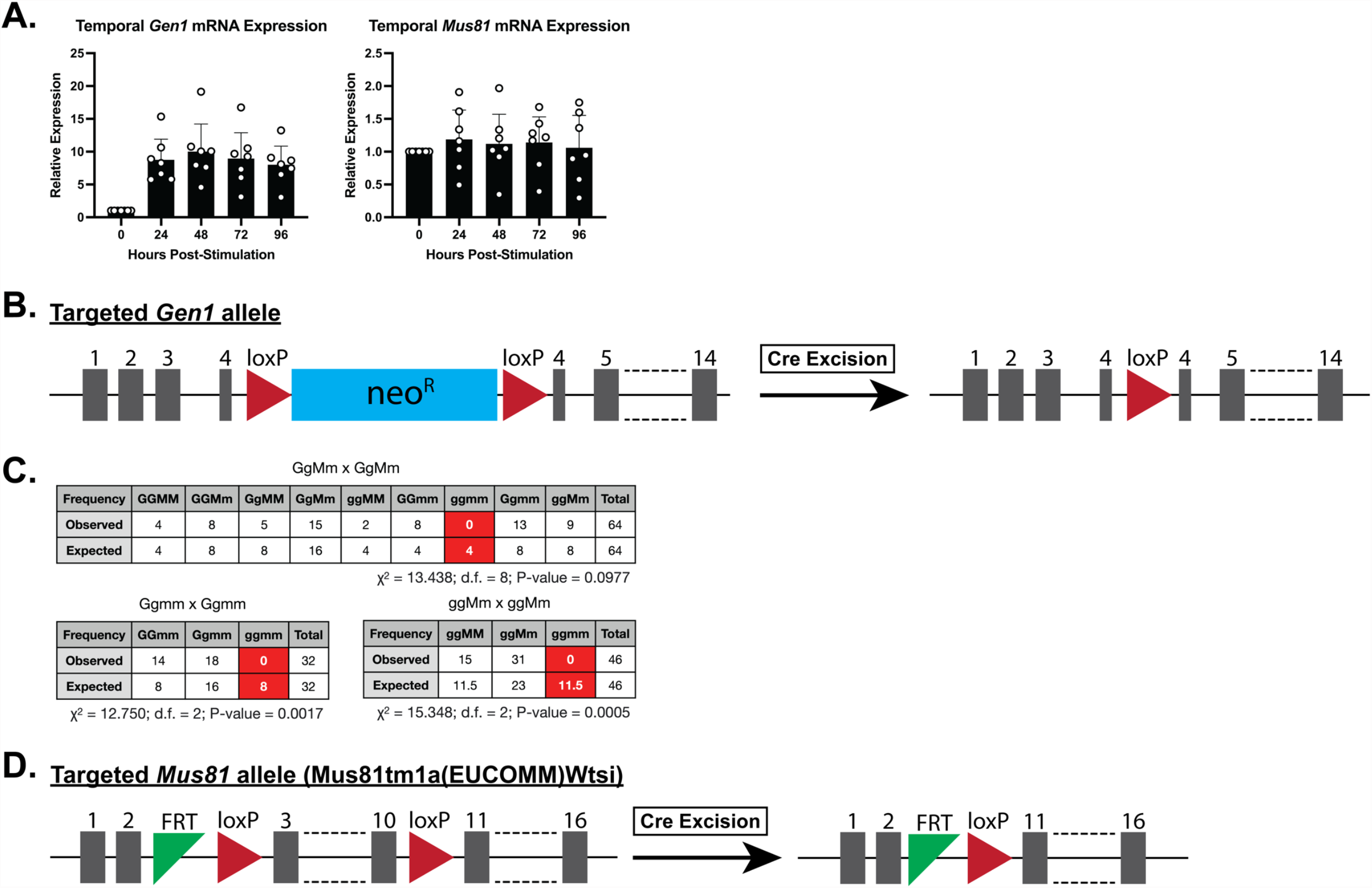
(A) Relative quantification of *Gen1* and *Mus81* mRNA transcript levels by RT-qPCR in wild-type mouse B cells stimulated in LPS+IL-4 culture from 0 hour to 96 hours post-stimulation. The expression level of the transcripts was normalized to the 0-hour time point. (B) Schematic of gene-targeting in generating the constitutive *Gen1*^-/-^ mouse model. The reading frame of exon 4 is disrupted with a floxed neomycin-resistance gene cassette that was subsequently excised upon Cre recombinase expression. (C) Tables depicting the observed and expected frequencies of the genotypes of pups generated from the indicated breeding schemes: G and M represent the wild-type alleles of *Gen1* and *Mus81*, respectively, whereas g and m represent the null alleles of *Gen1* and *Mus81*; d.f. denotes degree(s) of freedom. (D) Schematic of the *Mus81*^fl/fl^ allele in which exons 3 to 10 are flanked by loxP sites. When Cre recombinase is expressed under a tissue-specific promoter, exons 3 through 10 are excised, generating a null allele. FRT site is a remnant of pair that previously flanked a lacZ reporter-neomycin selection cassette.

**Figure 2–figure supplement 1.**
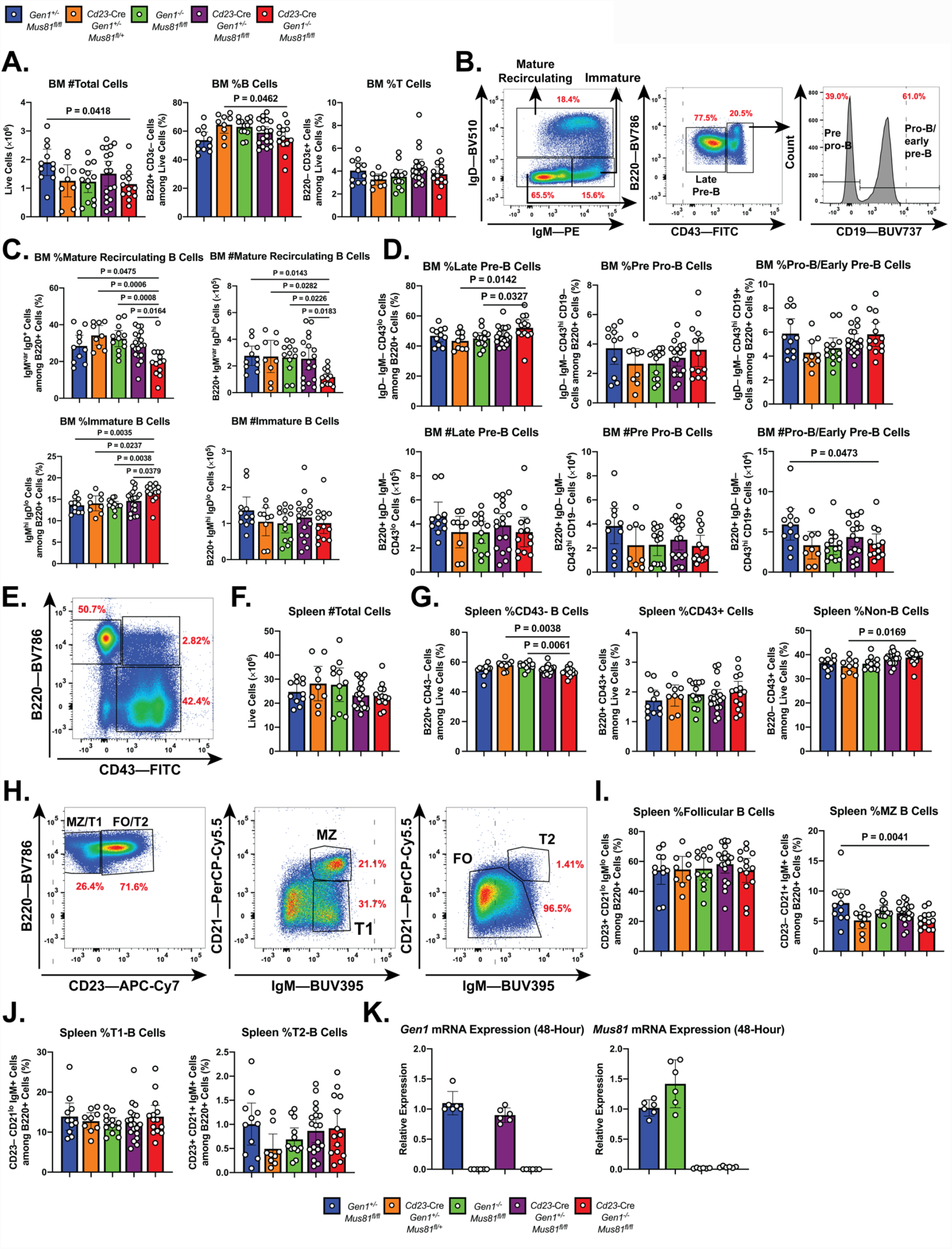
Steady-state phenotyping and quantification of the splenic and bone marrow B cell populations in the *Cd23*-Cre *Gen1*^-/-^ *Mus81*^fl/fl^ mice. A) Quantification of bone marrow cellularity and percentage of total B and T cell populations in the bone marrows of mice of the indicated genotypes. (B) Gating strategy of the B-cell populations in the bone marrow: mature recirculating, immature, late-pre-B, pre-pro-B, and pro-B/early pre-B cells. (C) Quantification of the percentage among live B220+ B cells and absolute counts of mature recirculating and immature B cells. (D and E) Quantification of frequencies (D) and absolute counts (E) of late pre-B, pre-pro-B, and pro-B/early pre-B cells among live B220+ B cells. (F) Representative flow cytometric plot depicting gating strategy of B220+ CD43– B, B220+ CD43+ B, and non-B (B220– CD43+) cells in the spleen. (G and H) Quantification of the absolute number of live splenocytes, and of the percentage of splenic CD43– B, CD43+ B, and non-B cells in the mice of the indicated genotypes. (I) Gating strategy of the various splenic B cell subpopulations: marginal zone (MZ), follicular (FO), transitional-1 (T1), and transitional-2 B cells. (J) Frequencies of follicular and marginal zone B cells among total splenic B220+ cells. (K) Frequencies of T1 and T2 B cells among whole B220+ B cells in the spleen. (L) Relative expression of *Gen1* and *Mus81* mRNA transcripts in LPS+IL-4-activated B cells of the indicated genotypes at 48 hours after stimulation. Data in (A)–(L) are from three independent experiments with 11 to 18 mice per genotype. Data in (M) are from three independent experiments with 6 mice per genotype. Bars depict the arithmetic mean and error bars represent the 95% confidence interval of the measured parameters. P-values were calculated with ordinary one-way ANOVA analysis with Dunnett’s multiple comparisons test without pairing. All means were compared to the *Cd23*-Cre *Gen1*^-/-^ *Mus81*^fl/fl^ group.

**Figure 3–figure supplement 1.**
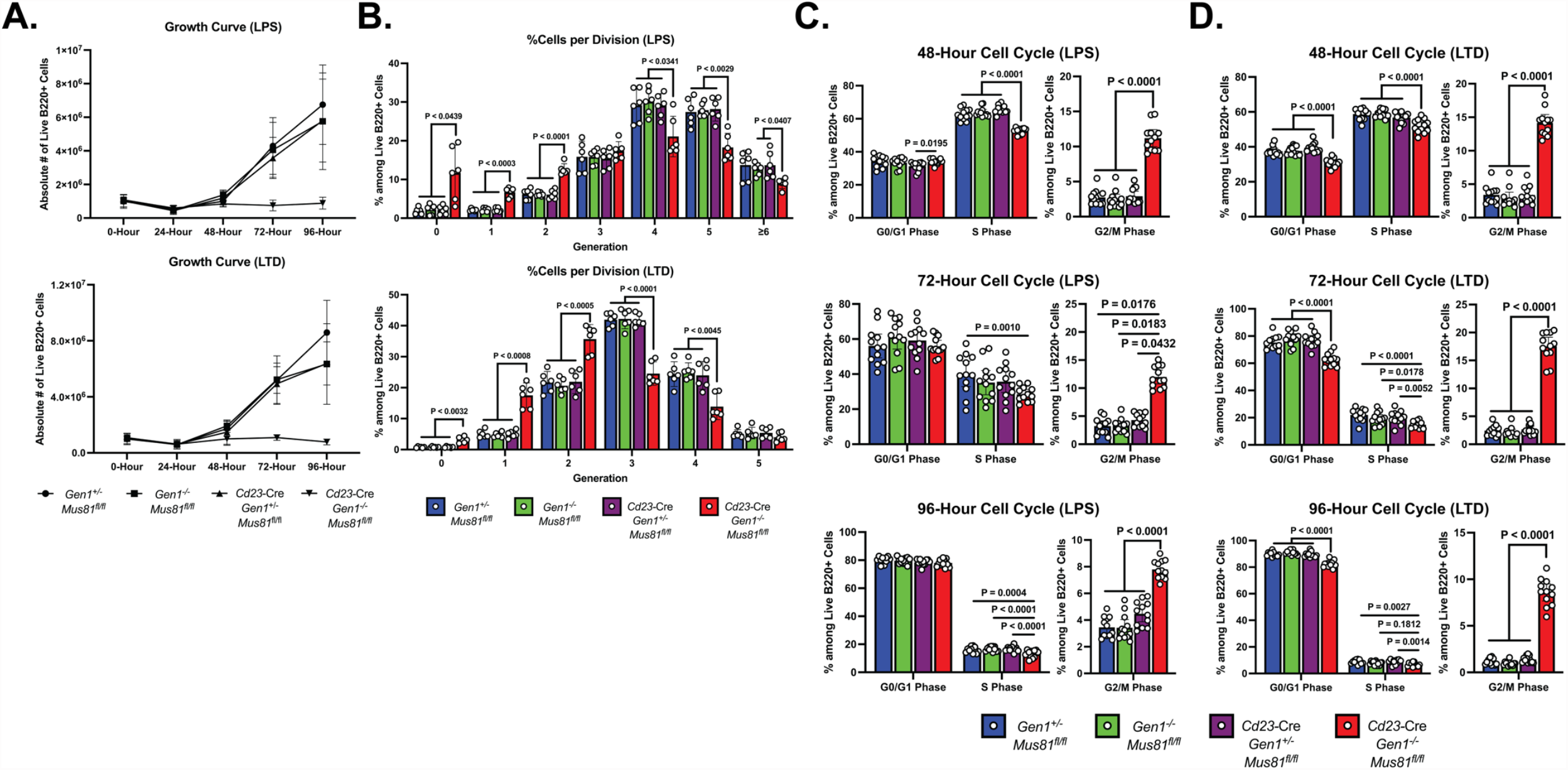
*Ex vivo* characterization of DKO cells following LPS and LPS+TGF-β+anti-IgD-dextran (LTD) stimulations. (A) Growth curve of LPS- and LTD-activated B cells. (B) Percentage of live B220+ cells in each generation. (C and D) Cell cycle analysis of *ex vivo*-activated B cells. Frequency of cells among live B220+ B cells in G0/G1, S, and G2/M phases at 48, 72, and 96 hours of LPS (C) or LTD (D) culture. Data in (A) are from four independent experiments with 9 mice per genotype. Data in (B) are from three independent experiments with 6 mice per genotype. Data in (C) and (D) are from six independent experiments with 12 mice per genotype. Bars display the arithmetic mean and error bars represent the 95% confidence interval of the measured parameters. P-values were determined using ordinary two-way ANOVA analysis with Dunnett’s multiple comparisons test without pairing wherein the mean of the *Cd23*-Cre *Gen1*^-/-^ *Mus81*^fl/fl^ group was compared to the rest.

**Figure 3–figure supplement 2.**
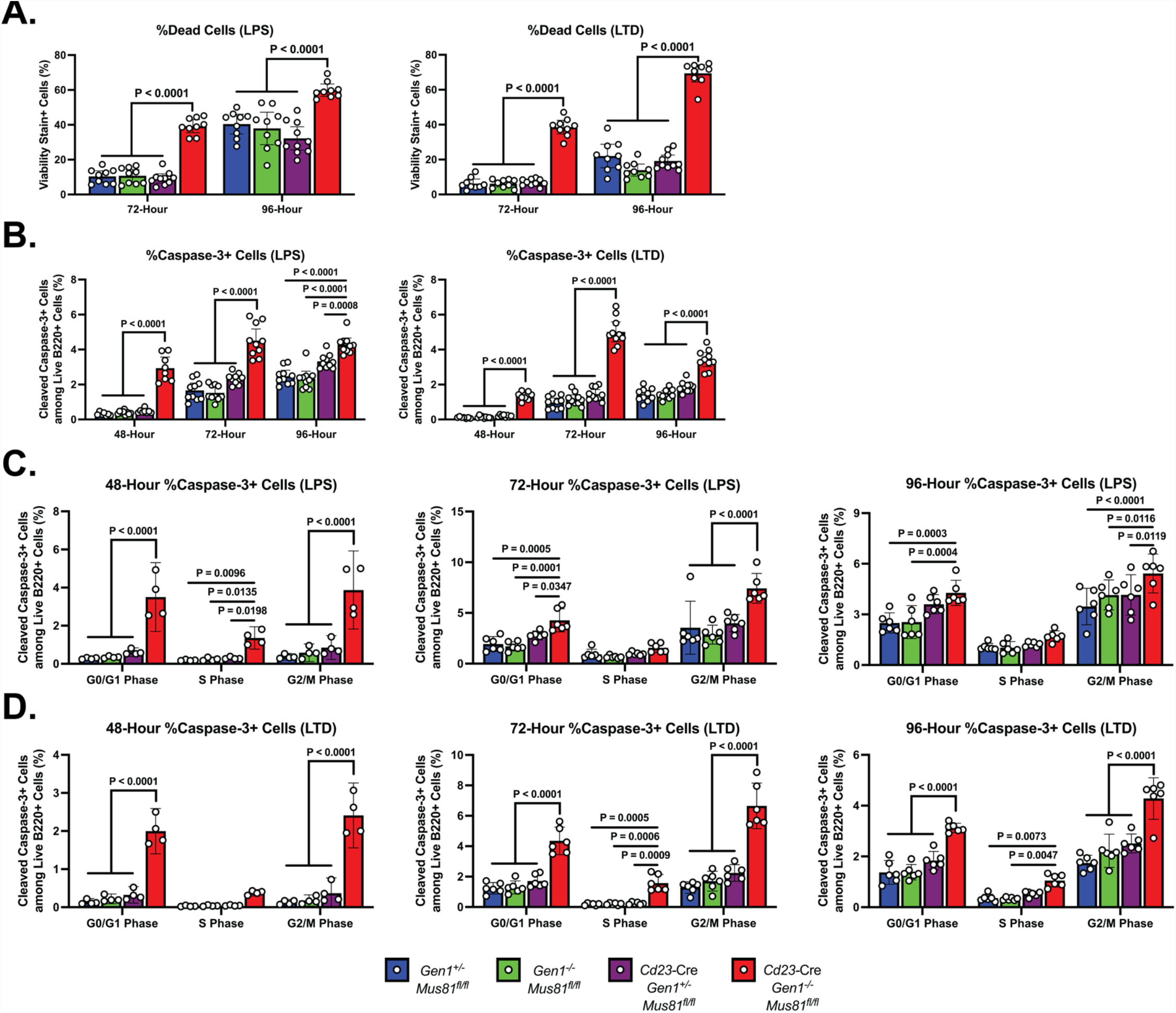
Cell death profiles of DKO cells stimulated in LPS and LTD cultures. (A) Frequency of dead cells quantified as a percentage of B220+ singlets staining for the viability dye after 72 and 96 hours of culture in media with LPS or LPS+TGF-β+anti-IgD-dextran (LTD). (B) Apoptosis in *ex vivo* B cell cultures. Percentage of cleaved caspase-3+ cells within the live B cell population at 48, 72, and 96 hours post-stimulation. (C and D) Cell cycle-specific determination of apoptosis in *ex vivo*-activated B cells. Frequency of cleaved caspase-3+ cells among live B220+ cells in each phase at 48, 72, and 96 hours of LPS (C) or LTD (D) culture. Data in (A) are from four independent experiments with 9 mice per genotype. Data in (B) are from five independent experiments with 8 to 10 mice per genotype. Data in (C) and (D) are from two (48-hour) or three (72- and 96-hour) independent experiments with 4 to 6 mice per genotype. Bars represent the arithmetic mean and the error bars depict the 95% confidence interval of the measured parameters. P-values were calculated using ordinary two-way ANOVA analysis with Dunnett’s multiple comparisons test in which the means were compared to that of the *Cd23*-Cre *Gen1*^-/-^ *Mus81*^fl/fl^ cohort.

**Figure 4–figure supplement 1.**
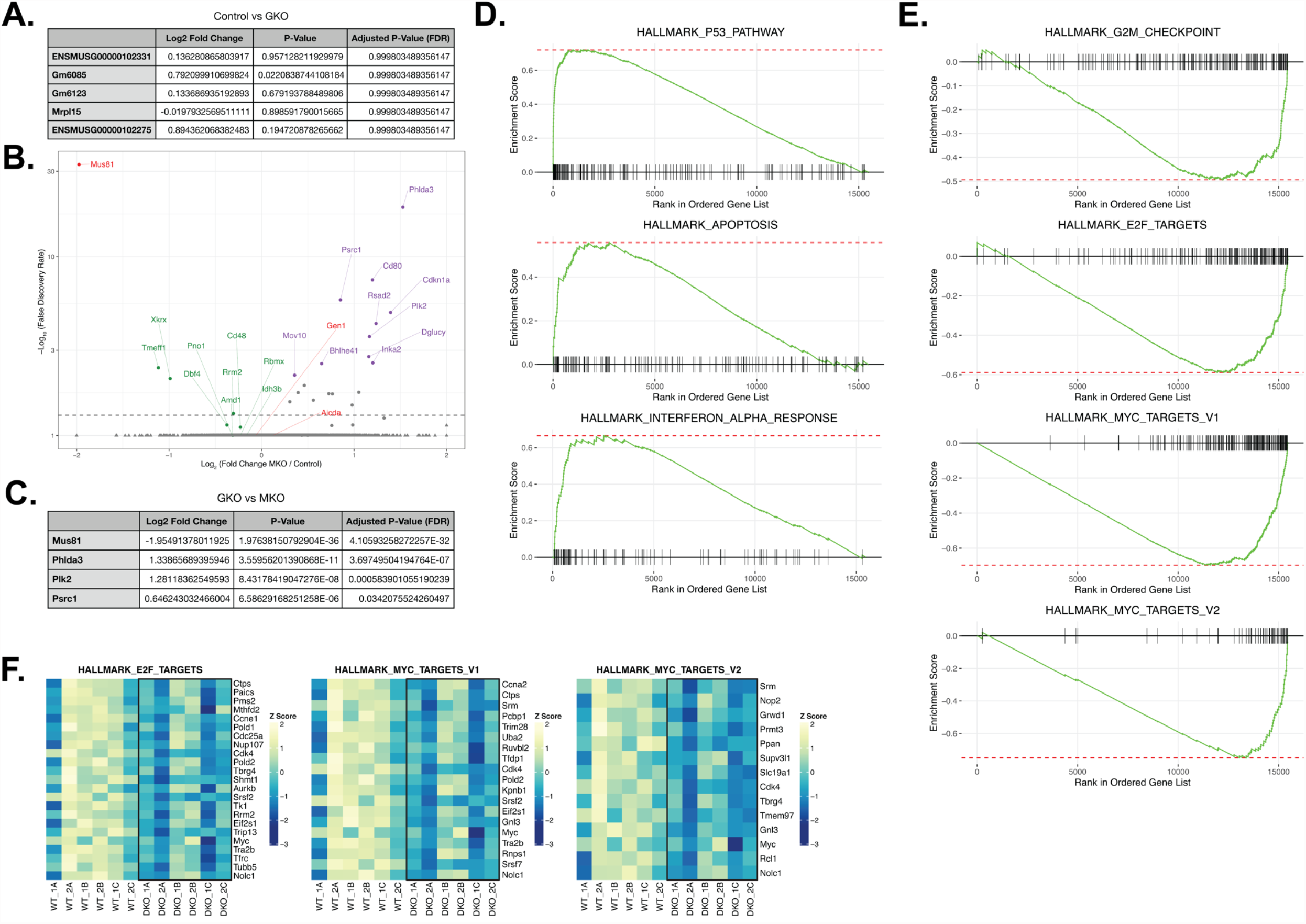
Transcriptomics and GSEA analyses of *ex vivo*-activated control, GKO, and MKO B cells. (A) Table listing the first five genes and the corresponding log_2_ fold change, P-value, and FDR generated from the DESeq analysis comparing *Gen1*^+/-^ *Mus81*^fl/fl^ (control) and *Gen1*^-/-^ *Mus81*^fl/fl^ (GKO) B cells (B) Volcano plot displaying the differential gene expression between control and *Cd23*-Cre *Gen1*^+/-^ *Mus81*^fl/fl^ (MKO) cultures at 48 hours post-stimulation. Purple dots indicate the top 10 significantly upregulated genes, and the green dots indicate genes that are de-enriched. (C) Table listing the genes that are differentially expressed between GKO and MKO B cells (D) GSEA plots of the Hallmark Gene Sets enriched in DKO cells. (E) GSEA plots of the Hallmark Gene Sets de-enriched in DKO cells. (F) Heatmap depicting the genes within the indicated Hallmark Gene Sets that had a log_2_ fold change of ≤–0.3 and FDR of <0.05 in DKO cells relative to control. Data are from 6 mice per genotype. The labels ‘1’ and ‘2’ represent male and female mice, respectively, and ‘A’ to ‘C’ denote the three experimental *ex vivo* groups.

